# GABAergic signaling in human and murine NK cells upon challenge with *Toxoplasma gondii*

**DOI:** 10.1101/2021.03.19.435611

**Authors:** Amol K. Bhandage, Laura M. Friedrich, Sachie Kanatani, Simon Jakobsson-Björkén, J. Ignacio Escrig-Larena, Arnika K. Wagner, Benedict J. Chambers, Antonio Barragan

**Author notes:** **Corresponding author**: Antonio Barragan, Dept. of Molecular Biosciences (MBW), The Wenner-Gren Institute, Stockholm University, Svante Arrhenius väg 20C, SE-106 91 Stockholm, Sweden.

## Abstract

Protective cytotoxic and proinflammatory cytokine responses by natural killer (NK) cells impact the outcome of infections by *Toxoplasma gondii*, a common parasite in humans and other vertebrates. However, *T. gondii* can also sequester within NK cells and downmodulate their effector functions. Recently, the implication of γ–aminobutyric acid (GABA) signaling in infection and inflammation-related responses of mononuclear phagocytes and T cells has become evident. Yet, the role of GABAergic signaling in NK cells has remained unknown. Here, we report that human and murine NK cells synthesize and secrete GABA in response to infection challenge. Parasitized NK cells secreted GABA while activation stimuli, such as IL-12/IL-18 or parasite lysates, failed to induce GABA secretion. GABA secretion by NK cells was associated to a transcriptional upregulation of GABA synthesis enzymes (GAD65/67) and was abrogated by GAD-inhibition. Further, NK cells expressed GABA-A receptor subunits and GABA signaling regulators, with transcriptional modulations taking place upon challenge with *T. gondii*. Exogenous GABA and GABA-containing supernatants from parasitized dendritic cells (DCs) impacted NK cell function by reducing the degranulation and cytotoxicity of NK cells. Conversely, GABA-containing supernatants from NK cells enhanced the migratory responses of parasitized DCs. This enhanced DC migration was abolished by GABA-A receptor antagonism or GAD-inhibition and was reconstituted by exogenous GABA. Jointly, the data show that NK cells are GABAergic cells and that GABA hampers NK cell cytotoxicity *in vitro*. We hypothesize that GABA secreted by parasitized immune cells modulates the immune responses to *T. gondii* infection.

**Summary sentence:** In response to infection challenge, NK cells synthesize and secrete *γ*-aminobutyric acid (GABA), which impacts NK cell and dendritic cell functions via GABA-A receptors.

## 1 Introduction

G–aminobutyric acid (GABA), the main inhibitory neurotransmitter of the vertebrate brain, has also been attributed motogenic functions outside the central nervous system, including immune cell migration in infection and metastasis of cancer cells ^1-4^. GABA-A receptors (GABA-A R) are ionotropic chloride (Cl^-^) channels composed from pentameric combinations of 19 different subunits ^5^ and whose GABA signaling functions are regulated by cation-chloride co-transporters (CCCs) ^6^. GABAergic cells, for example neurons, synthesize GABA via glutamate decarboxylases (GAD65/67) and metabolize GABA by GABA-transaminase (GABA-T) ^7^. Further, GABA transporters (GAT) shuttle GABA in and out of cells ^8^.

The roles of GABAergic signaling have been extensively studied in neurons and astrocytes ^9,10^ but remain chiefly unexplored in cells of the immune system ^11^. Recent reports have elucidated that immune cells, such as T cells, macrophages, monocytes and dendritic cells (DCs), express GABAergic signaling components and are responsive to GABA, which has been attributed immunomodulatory functions ^2,3,12,13^. However, GABAergic signaling in natural killer (NK) cells has remained elusive.

*Toxoplasma gondii* is an obligate intracellular protozoan that infects warm-blooded vertebrates, including humans and rodents ^14^. Chronic carriage of *T. gondii* without major or no symptomatology is common. However, systemic dissemination of *T. gondii* can cause life-threatening encephalitis in immune-compromised patients, disabling disease in the developing foetus, and ocular manifestations ^15,16^. The tachyzoite parasite stage actively invades and replicates within nucleated cells in the host ^17^, including immune cells such as dendritic cells (DCs), T cells or NK cells ^18-20^. Upon infection of DCs and other mononuclear phagocytes, *T. gondii* promotes migratory activation of parasitized phagocytes through GABAergic signaling to facilitate parasite dissemination via a *Trojan horse* mechanism ^3,18,21^.

NK cells play important roles in *T. gondii* infection, with an impact on innate, adaptive and regulative responses ^22^. Early studies showed that NK cells, alike DCs, have a protective role in *T. gondii* infection as they serve as an early source of Interleukin-12 (IL-12) and Interferon-γ (IFN-γ) ^23-25^. IFN-γ responses lead to the activation and differentiation of macrophages and DCs which enhances the killing of the parasite and supports the activation of the T cell responses ^26,27^. In this setting, NK cells home to lymphoid organs where they interact with phagocytes ^28^, display enhanced motility ^29^ and are able to kill *T. gondii* -infected target cells *in vitro* and in vivo ^20^. Interestingly, NK cell-mediated killing of parasitized DCs led to rapid egress of viable parasites which in turn infected the effector NK cells ^20^. Conversely, parasitized NK cells exhibited impaired recognition of target cells and reduced cytokine release ^30^.

Yet, the host-pathogen interplay between *T. gondii* and parasitized NK cells and DCs remains elusive. Here, we report that human and murine NK cells express the components of a GABAergic machinery and secrete GABA upon challenge with *T. gondii* tachyzoites. We test the functional implications of GABA secretion on DC migration and effector functions of NK cells.

## 2 Methods

### 2.1 Ethics statement

The Regional Ethics Committee, Stockholm, Sweden, approved protocols involving human cells. All donors received written and oral information upon donation of blood at the Karolinska University Hospital. All the animal experimentation procedures involving infection and extraction of cells/organs from mice were approved by Regional Animal Research Ethical Board, Stockholm, Sweden in concordance with in EU legislation (permit numbers 9707/2018 and 14458/2019).

### 2.2 Experimental animals

Six to 10-week-old C57BL/6NCrl mice were purchased from Charles River (Sulzfeld, Germany) and maintained or bred under pathogen-free conditions at Experimental Core Facility (ECF), Stockholm University, Sweden. Six to 10-week-old B6.RAG1^-/-^ mice ^31^ and C57BL6/J mice (Janvier Laboratories, France) were maintained at Astrid Fagraeus Laboratories, Karolinska Institutet.

### 2.3 Parasites and cell lines

*T. gondii* GFP-expressing RH-LDM (type I) and ME49/PTG (type II) ^32,33^ were maintained by serial 2-day passaging in human foreskin fibroblast (HFF-1, ATCC® SCRC-1041™, American Tissue Culture Collection) monolayers cultured in DMEM (Thermofisher scientific) with 10% fetal bovine serum (FBS; Sigma-Aldrich), gentamicin (20 µg/ml; Gibco), glutamine (2 mM; Gibco), and HEPES (0.01 M; Gibco). NK-92 cells (ATCC® CRL-2407™) were grown in stem cell growth medium (CellGro; CellGenix, Freiburg, Germany) with 20% heat-inactivated FBS (Gibco, Life Technologies, Carlsbad, CA, USA) and 1000 U/mL of Proleukin (Novartis, Basel, Switzerland) or IL-2 (Miltenyi, Bergisch Gladbach, Germany). YAC-1 cells (ATCC® TIB-160™) and K-562 cells (ATCC® CCL-243™) were maintained in RPMI 1640; 10 mM HEPES, 2 × 10^−5^ M 2-mercapto-ethanol (2-ME), 10% FBS, 100 U/ml penicillin, 100 U/ml streptomycin with 500U/ml IL-2.

### 2.4 Primary cells

#### NK cells

Single-cell suspensions from spleens were depleted of erythrocytes, and NK cells were positively sorted using anti-DX5^+^ magnetic beads or by negative sorting using MACS separation, according to the manufacturer’s instructions (Miltenyi Biotec, Bergisch Gladbach, Germany). Cells were resuspended in RPMI 1640; 10 mM HEPES, 2 × 10^−5^ M 2-ME, 10% FBS, 100 U/ml penicillin, 100 U/ml streptomycin with 1000 U/ml mouse IL-2 (Immunotools). For mRNA expression by real-time quantitative PCR, NK cells were first isolated by negative sorting using NK cell isolation kit (Miltenyi Biotech) and further purified by flow cell sorting following labelling with anti-CD3 and anti-NK1.1 (FACS Aria Fusion, BD Biosciences). Cell isolations with purity ranging between 96,1-99,8% were used (Supplementary Fig. 1). Human NK cells were isolated from peripheral blood mononuclear cells using human NK cells isolation kit (Miltenyi Biotech). Cells were then incubated overnight in 100 U/ml IL-2 (Peprotech) overnight prior to being used in experiments. Human NK cells challenged with GFP-expressing *T. gondii* (RH-LDM) were sorted (FACS Aria Fusion, BD Biosciences) based on GFP-expression and CD56 expression. GFP^+^ cells (defined as infected) and GFP^-^ cells (defined as bystander) were collected and incubated over night before supernatants were collected and analyzed by GABA-ELISA.

#### DCs

Mouse bone marrow-derived DCs (mBMDCs) were generated as previously described ^3^. Briefly, bone marrow cells extracted from legs of 6-10-week-old C57BL/6 mice (Charles River) were cultivated in RPMI 1640 with 10% FBS, gentamicin, glutamine and HEPES, additionally supplemented with recombinant mouse GM-CSF (10 ng/ml; Peprotech) at 37°C. Medium was replenished on days 2 and 4. Loosely adherent cells harvested on day 6-8 were used for experiments.

### 2.5 Real-time quantitative PCR

Total RNAs were extracted using Direct-zol miniprep RNA kits (Zymo Research) with TRI reagent (Sigma-Aldrich) and first-strand cDNA was synthesized using Superscript IV (Invitrogen) using a standard protocol. Real-time quantitative PCR (qPCR) was performed in QuantStudio 5 384 Optical well plate system (Applied Biosystem) in a standard 10 ul with the 2X SYBR FAST qPCR Master Mix (Sigma-Aldrich) with gene specific primers (**Supplementary Table 1**) with a standard amplification and melt curve protocol. When indicated, primers were validated on mouse whole brain homogenates, prepared as previously described. ^34^. Relative expression (2^-ΔCt^) was determined for each target in relation to a normalization factor, geometric mean of reference genes, TATA-binding protein (TBP) and importin 8 (IPO8). Heat maps represent transcriptional changes in mRNA expression (2^-ΔCt^) upon challenge with *T. gondii* tachyzoites (RH-LDM).

### 2.6 GABA enzyme-linked immunosorbent assays

NK cells were plated at a density of 1×10^6^ cells/ml and challenged with freshly-egressed *T. gondii* tachyzoites (RH-LDM, MOI 2, 24 h) in presence or absence of GABA synthesis inhibitor semicarbazide (SC, 50 μM, Sigma-Aldrich). Cells were also challenged for 24 h with MOI-equivalent numbers of heat-inactivated (HI, at 56°C for 30 min) tachyzoites or sonicated tachyzoites (30 s at 10 microns; Soniprep 150, MSE), supernatants released by tachyzoites in 1 h or, in presence of recombinant mouse IL-12 (100 ng/ml, Peprotech, Rocky Hill, NJ) and recombinant mouse IL-18 (100 ng/ml, Biosource, Brussels, Belgium). GABA concentrations in supernatants were quantified from the standard curve generated at a wavelength of 450 nm (VMax® Kinetic ELISA Microplate Reader, Molecular Devices) by ELISA (Labor Diagnostica Nord, Nordhorn, Germany) as described previously ^3^.

### 2.7 Motility assays

Cell motility analyses was performed as previously described ^3,35^. Briefly, mouse bone marrow-derived DCs were challenged with freshly-egressed ME49/PTG (MOI 3, 4-6 h) tachyzoites. Combined treatments were performed, as indicated in figure legend, with GABA synthesis inhibitor (SC, 50 μM), supernatants from T. gondii-challenged non-treated or SC-treated mNK cells (added at medium:supernatant ratio of 1:1), GABA-A R inhibitor picrotoxin (50 μM, Sigma-Aldrich) or GABA (5 μM, Sigma-Aldrich). Cells were then imaged in 96-well plates every min for 60-90 min (Zeiss Observer Z.1). Motility tracks for 50-60 cells per treatment were analyzed for each experiment by manual tracking in ImageJ software. Motility plots were generated with chemotaxis and migration stand-alone tool (version 2.0, IBIDI). X-and y-axes in the plots show distances in μm. The Box-and-whisker dot plots represent median velocities (μm/min) with boxes marking 25^th^ to 75^th^ percentile and whiskers marking 10^th^ and 90^th^ percentiles of the datasets. Grey circles represent velocities from individual cells.

### 2.8 NK cell degranulation assay

**Murine NK cells** were incubated for 2 h in the presence of GABA (2 or 10 μM, Sigma-Aldrich), supernatants from unchallenged mBMDCs or supernatants from mBMDCs challenged with *T. gondii* (PTG, MOI2, 24 h). Supernatants were added at a medium:supernatant volume ratio of 1:1. NK cells were then washed and co-cultured with YAC-1 cells at a ratio of 10:1 in the presence of anti-CD107a antibody (LAMP-1, eBioscience) diluted 1:200) The cells were briefly centrifuged and after 30 min, monensin (BioLegend) was added to the culture for further 1.5 h. After this time point, cells were stained for CD3 (clone 145-2C11, Biolegend), NK1.1 (PK136, Biolegend) and fixable viability dye (eFluor 780, eBioscience). Flow cytometry was performed on CyAN ADP LX 9-colour flow cytometer (Beckman Coulter, Pasadena, CA).

### 2.9 Cytotoxicity assay

**Human NK-92 cells** were preincubated with GABA for 2 h prior to use in the cytotoxicity assay and washed prior to use in the cytotoxicity assay. K-562 cells were incubated for 1 h in the presence of Na_2_ ^51^CrO_4_ (^51^Cr, Perkin Elmer), washed twice, and incubated with NK-92 cells at the indicated effector:target (E:T) ratios. After 2 h, cell culture supernatants were collected and analyzed by a radiation counter (Wallac, PerkinElmer). Specific lysis was calculated as follows: % specific lysis [(experimental release x spontaneous release) / (maximum release x spontaneous release)] x100.

### 2.10 Western blot

Human NK cells were challenged with freshly-egressed *T. gondii* tachyzoites (RH-LDM, MOI 2, 24 h). Cells were harvested and lysed in RIPA buffer with protease inhibitor cocktail (Roche), sonicated, diluted with 4X Laemmli buffer and boiled. Samples were subjected to SDS-PAGE on 8% polyacrylamide gels, transferred onto PVDF membrane (Hybond P, Merck Millipore), blocked in 5% BSA for 30 min and incubated ON with mouse anti-GAD67 monoclonal antibody (1:6000, MAB5406, Merck Millipore) or 1 h with rabbit polyclonal anti-GAPDH antibody (1:3000, ABS16, Merck Millipore) followed by incubation with respective HRP-conjugated secondary antibodies (G21040, Thermofisher; 7074S, Cell signaling). Protein bands were revealed by enhanced chemiluminescence reagents (ECL™ Prime, Merck Millipore) in a ChemiDoc XRS+ imaging system (BioRad, Stockholm, Sweden). Mouse brain protein was used as positive control.

### 2.11 Statistical analyses

Data mining and statistical analyses were performed using GraphPad Prism 7.0 (La Jolla, CA, USA) and described in figure legends. Statistical significance was defined as p < 0.05.

### 2.12 Online Supplemental material

**Supplementary Table 1**. Primer pair sequences used in real-time quantitative PCR

**Supplementary Table 2**. Transcriptional expression of GABA-A R subunits, GABA enzymes, GABA transporters and CCCs in *T. gondii*-challenged primary mNK cells

**Supplementary Table 3**. Transcriptional expression of GABA-A R subunits, GABA enzymes, GABA transporters and CCCs in *T. gondii*-challenged primary hNK cells

**Supplementary Table 4**. Correlation analyses of mRNA expression levels with amounts of GABA released by *T. gondii*-challenged NK cells from individual human donors

**Supplementary Figure 1**. Representative flow cytometric analysis of cell purity of mNK cells used for mRNA analyses of GABA-related genes

**Supplementary Figure 2**. Relative mRNA expression of 19 GABA-A R subunits in NK cells from four different human donors.

## 3 Results

### 3.1 Murine and human NK cells transcribe a GABAergic machinery

Whether NK cells express a GABAergic machinery has remained elusive. We therefore assessed mouse and human NK cells for transcripts of GABA signaling components. Interestingly, both murine and human NK cells expressed a number of transcripts of GABA-A receptors (GABA-A R) subunits (**Fig. 1A, Supplementary Fig. 2**). Commonly expressed GABA-A R subunits for mouse and human NK cells were α3, β2-3, γ1, δ and ρ1-2, while additional subunits were not amplified or only expressed in mouse or in human NK cells (**Table 1**). Furthermore, NK cells expressed transcripts of the GABA synthesis enzymes glutamate decarboxylase (GAD65/67), GAD67 being the most prominently expressed, and also the degrading enzyme GABA transaminase (GABA-T) (**Fig. 1B**). Interestingly, among known GABA transporters, only transcripts of GAT2 were detected, similarly in mouse and human NK cells (**Fig. 1C**). Finally, a number of cation-chloride cotransporters (CCC), known to regulate GABA signaling, were transcribed in both cell types (**Fig. 1D**). We conclude that human and mouse NK cells transcribe a set of GABAergic signaling components that, in theory, suffice to synthesize, degrade and transport GABA, and form functional GABA-A Rs.

**Table 1.** GABA-A Receptor subunits transcribed by NK cells.

| Cell / tissue | Condition | $\alpha 1$ | $\alpha 2$ | $\alpha 3$ | $\alpha 4$ | $\alpha 5$ | $\alpha 6$ | $\beta 1$ | $\beta 2$ | $\beta 3$ | $\gamma 1$ | $\gamma 2$ | $\gamma 3$ | $\delta$ | $\varepsilon$ | $\theta$ | $\pi$ | $\rho 1$ | $\rho 2$ | $\rho 3$ |
| --- | --- | --- | --- | --- | --- | --- | --- | --- | --- | --- | --- | --- | --- | --- | --- | --- | --- | --- | --- | --- |
| mNK | Unchallenged |  | 6/8* | 3/8 |  |  |  |  | 6/8 | 7/8 | 7/8 | 4/8 |  | 8/8 | 7/8 |  | 3/8 | 8/8 | 4/8 |  |
|  | *T. gondii*-challenged |  | 5/8 | 3/8 |  |  |  |  | 7/8 | 8/8 | 7/8 | 4/8 |  | 8/8 | 8/8 |  | 3/8 | 8/8 | 7/8 |  |
| hNK | Unchallenged |  |  | 4/4 |  | 4/4 | 4/4 |  | 4/4 | 1/4 | 3/4 |  |  | 3/4 |  | 3/4 |  | 3/4 | 4/4 |  |
|  | *T. gondii*-challenged |  |  | 4/4 |  | 4/4 | 4/4 |  | 4/4 | 1/4 | 4/4 |  |  | 3/4 |  | 3/4 |  | 4/4 | 4/4 |  |
| Mouse brain | Unchallenged | 4/4 | 4/4 | 4/4 | 4/4 | 4/4 | 4/4 | 4/4 | 4/4 | 4/4 | 4/4 | 4/4 | 4/4 | 4/4 | 4/4 | 4/4 | 4/4 | 4/4 | 4/4 | 4/4 |

Murine NK cells (mNK) and human NK cells (hNK) were assessed for mRNA expression of GABA-A receptor subunits after challenge with *T. gondii* as detailed under Methods. Mouse whole brain homogenate mRNA was used as reference material.
\* indicates number of mice or human donors, respectively, expressing a particular subunit related to the total number of independent samples analyzed (mNK, n = 8 mice; hNK, n = 4 human donors; mouse brain homogenate, n = 4 mice).
Blank spaces indicate that the particular subunit mRNA was not amplified in any sample.

**Figure 1.**
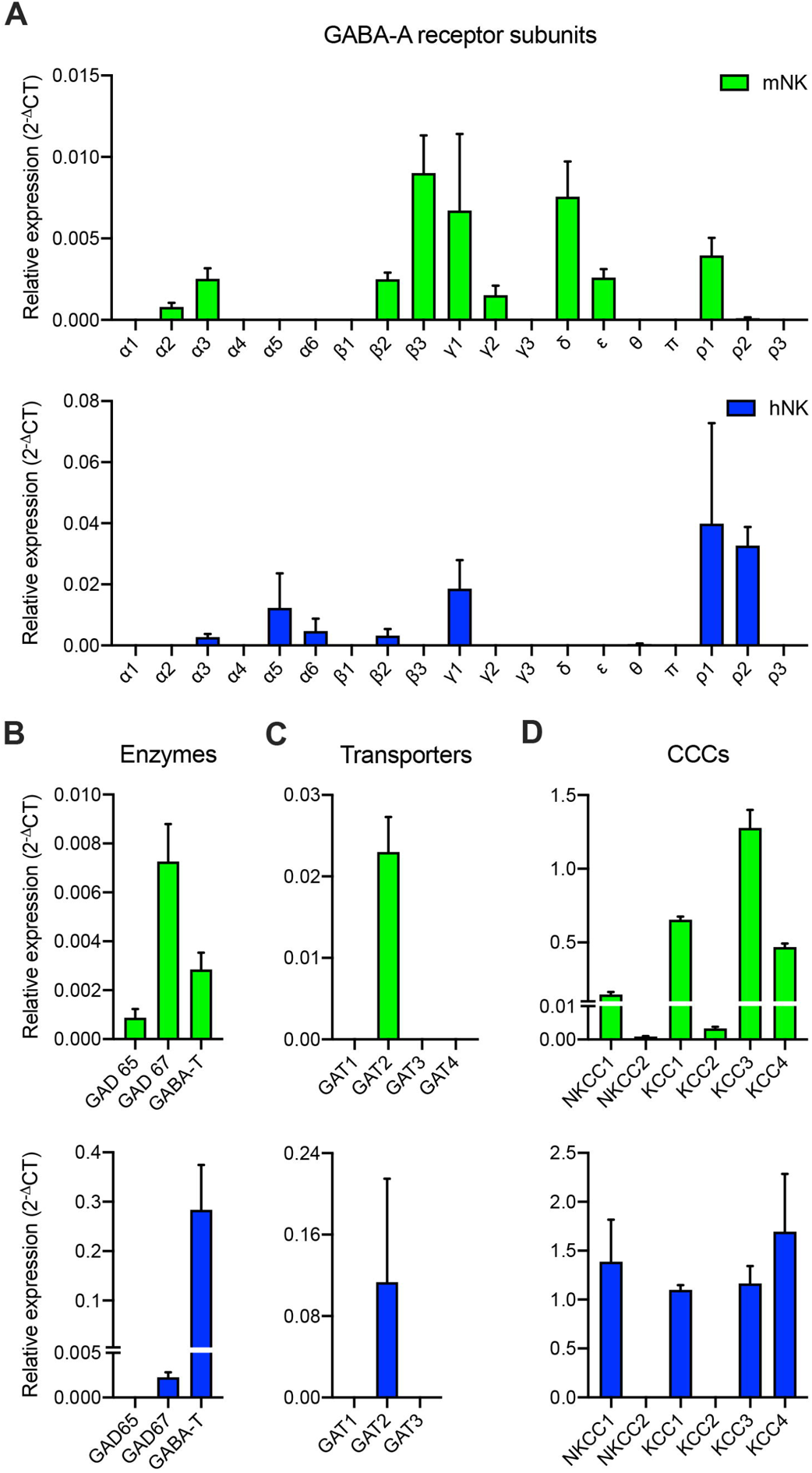
Expression of GABA signaling components by murine and human NK cells. Bar graphs show relative mRNA expression (2^-ΔCt^ +SEM) of **(A)** GABA-A R subunits, **(B)** GABA synthesis and catabolic enzymes (GAD65/67 and GABA-T, respectively), **(C)** GABA transporters and **(D)** cation-chloride co-transporters (CCCs) in mouse NK cells (mNK, n = 8 mice) and human NK cells (hNK, n = 4 human donors).

### 3.2 Transcriptional modulation of GABAergic components in NK cells upon challenge with *T. gondii*

Next, we addressed if challenge of NK cells with *T. gondii* tachyzoites impacted the transcriptional expression of GABAergic signaling components. We previously reported that *T. gondii* actively invades and replicates within NK cells ^20^. Overall, a transcriptional upregulation was observed for multiple GABAergic components (**Fig. 2, Supplementary Tables 2 and 3**). Interestingly, up and downregulations were observed for GABA-A R subunits (**Fig. 2A**) and the GABA regulators CCCs (**Fig. 2D**), indicating a significant impact of infection on their expression. In contrast, the transcription of GABA synthesis enzymes (**Fig. 2B**) and transporters (**Fig. 2C**) were consistently upregulated, with the exception of the GABA degrading enzyme GABA-T in human NK cells. We conclude that infection challenge led to upregulated expression of multiple GABAergic signaling components in NK cells.

**Figure 2.**
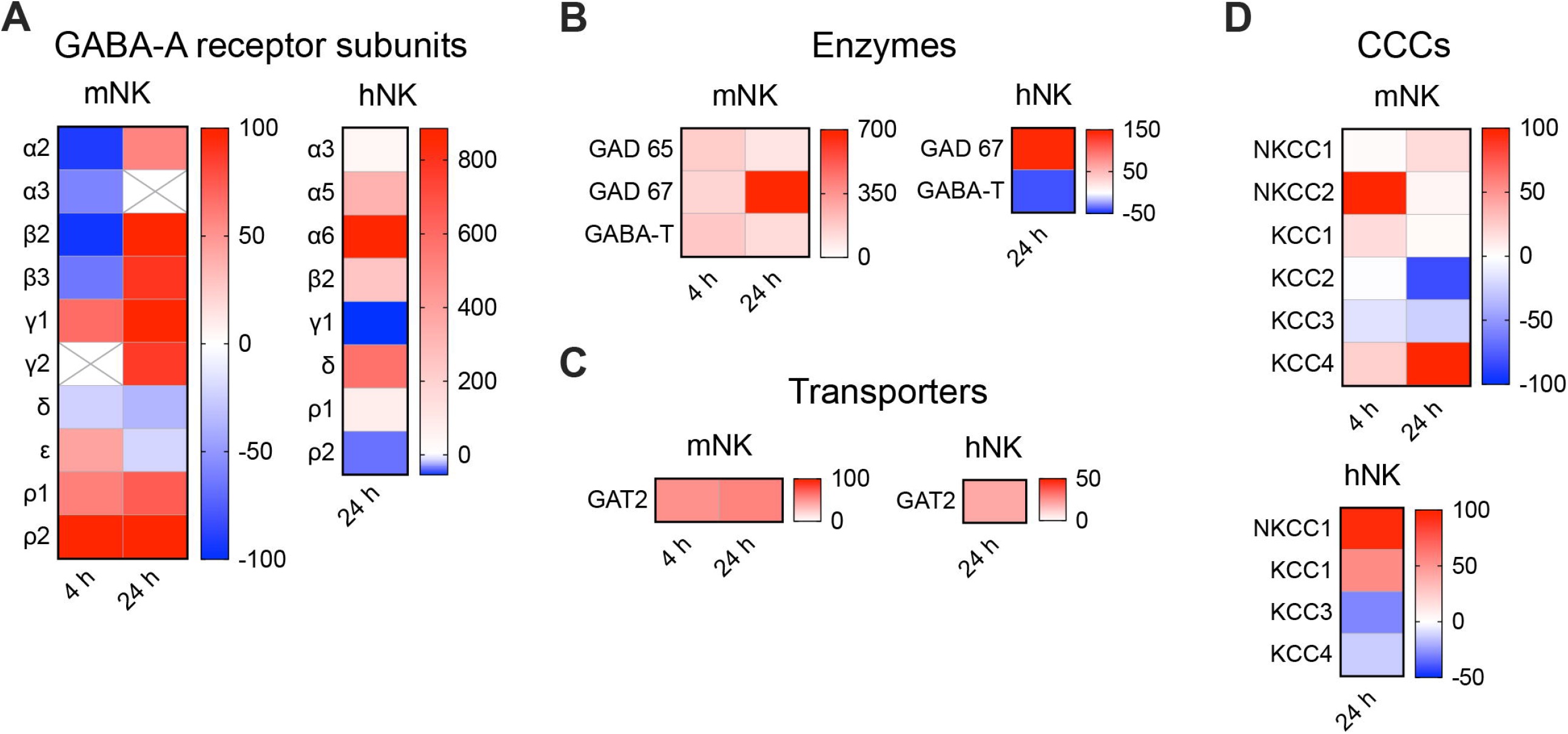
Transcriptional modulation in GABA signaling components in NK cells upon *T. gondii* challenge. **(A-D)** Heat maps show transcriptional changes of indicated GABA signaling components upon challenge of murine NK cells (mNK) and human NK cells (hNK) with *T. gondii* (RH-LDM), at indicated time points. Red and blue color scales indicate, for each gene, the percentage increase or decrease in expression, respectively, normalized to unchallenged cells at the same time point. (X) indicates no amplification. Detailed numeric analyses of murine and human NK cells are provided in Supplementary Tables 2 and 3, respectively (n=3-5 independent experiments).

### 3.3 *T. gondii*-infection induces GABA synthesis and secretion by human and murine NK cells

Next, we sought to determine if NK cells produce and secrete GABA. First, in *T. gondii*-challenged NK cells from human donors, we confirmed the presence of protein bands corresponding to the synthesis enzyme GAD67 from mouse brain homogenates (**Fig. 3A**), consistent with the observed transcriptional upregulation of GAD67 mRNA (**Fig. 2B**). Importantly, upon challenge with *T. gondii*, GABA-ELISA analyses revealed a dramatic elevation of GABA in the supernatant for both mouse and human NK cells (**Fig. 3B, C**). In presence of the GAD-inhibitor semicarbazide (SC), the amounts of GABA in the supernatant were significantly reduced (**Fig. 3D**). Moreover, MOI-equivalent doses of heat-inactivated tachyzoites, tachyzoite lysates and supernatants or addition of cytokines failed to induce GABA secretion (**Fig. 3D**), indicating that live intercellular parasites were necessary for GABAergic activation. Next, we explored GABA responses by individual human donors. NK cells from 9 human donors consistently responded to *T. gondii* challenge with elevated GABA secretion, and with variations among donors in the amounts of secreted GABA (Fig. 3E). This variation motivated an analysis of the expression of the GABA synthesis enzyme GAD67 and the GABA catabolic enzyme GABA-T for each donor. Interestingly, the amounts of secreted GABA by the individual donors were strongly correlated with an upregulation of GAD67 and downregulation of GABA-T expression, respectively (**Fig. 3F**). This correlation was absent for GABAergic genes unrelated to GABA metabolism (**Supplementary Table 4**). Finally, *T. gondii*-challenged NK cells were sorted in infected cells and by-stander cells (**Fig. 3G**). Supernatants from sorted infected NK cells presented significantly elevated GABA production related to supernatants from by-stander cells (**Fig 3H**). However, GABA concentrations were elevated in supernatants from by-stander cells related to supernatants from unchallenged cells (**Fig 3H**), indicating a minor by-stander effect, and with variations among human donors (**Fig 3I**). Jointly, the data demonstrate that NK cells synthesize and secrete GABA, which is dramatically accentuated upon *T. gondii* infection. Moreover, the data are indicative that the elevated amounts of secreted GABA by *T. gondii*-infected NK cells are mediated by increased GABA synthesis (GAD67) and decreased GABA catabolism (GABA-T).

**Figure 3.**
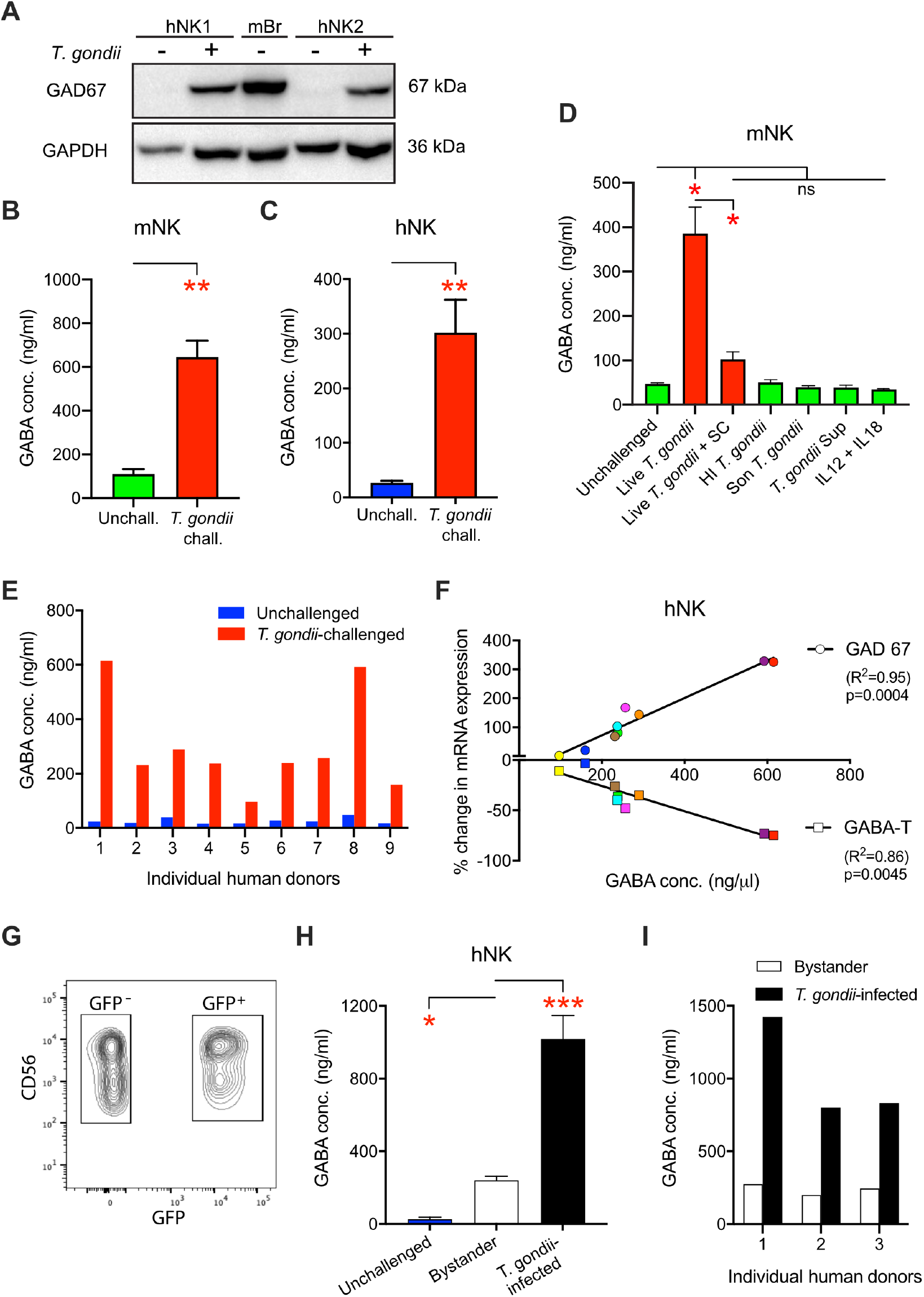
GABA secretion by human and murine NK cells. **(A)** Representative Western blot of lysates from unchallenged and T. gondii-challenged NK cells from 2 human donors (hNK1 and 2, respectively), immunoblotted with GAD67 antibody as indicated under Methods. Mouse brain lysate (mBr) served as a positive control and GAPDH immunoblotting was performed for loading reference. **(B, C)** Bar graphs show concentrations (mean +SEM) of GABA secreted in supernatants of (B) mouse NK cells (mNK, n = 6 mice) and (C) human NK cells (hNK, n = 9 human donors), respectively, challenged with *T. gondii* (RH-LDM) and quantified by ELISA. ** p < 0.01, Mann-Whitney test. (D) GABA concentration in mNK supernatants after 24 h of challenge with *T. gondii* (MOI 2) in presence and absence of GABA synthesis inhibitor (SC), or with MOI-equivalent amounts of heat-inactivated (HI) T. gondii, lysate (sonicated T. gondii), supernatants from T. gondii, or cytokine stimulation with IL-12 and IL-18, as detailed in Methods. ns: non-significant, * *p* < 0.05, one-way ANOVA followed by Tukey multiple comparison test. **(E)** GABA concentrations upon T. gondii-challenge of NK cells from individual human donors (1-9). **(F)** Correlation analyses of transcriptional expression changes of GABA metabolic enzymes with measured GABA concentrations in supernatants, for 9 human donors. For each donor, GABA concentration in the supernatants of hNK cells was correlated with transcriptional changes in GAD67 and GABA-T upon *T. gondii* challenge. Correlation was assessed by non-parametric spearman test. R^2^ indicates the linearity of the trend line. **(G)** Bivariate contour plots of hNK cells challenged with GFP-expressing *T. gondii* (RH-LDM) and stained for CD56. GFP^+^ CD56^+^ cells were defined as infected NK cells and GFP^-^ CD56^+^ cells as by-stander NK cells. **(H)** Bar graph shows GABA concentrations in spernatants from cell-sorted unchallenged, by-stander and infected hNK cells. * *p* < 0.05, *** *p* < 0.001, one-way ANOVA followed by Dunnett’s multiple comparisons test (n = 3 human donors). **(I)** GABA concentrations in supernatants from cell-sorted by-stander and infected hNK cells from 3 human donors (1-3).

### 3.4 Exogenous GABA and supernatants from *T. gondii*-infected DCs hamper NK cell degranulation and cytotoxicity

Previously, we found that murine NK cells infected with *T. gondii* exhibit impaired cytotoxicity towards YAC-1, the prototypic NK cell target cell ^30^. To test if GABA impacted this central effector function of NK cells, degranulation and cytotoxicity assays were performed in the presence of exogenous GABA at a concentration range corresponding to that attained in supernatants of DCs challenged with *T. gondii* ^3^ and challenged NK cells (low μM range) (Fig. 3). In presence of GABA, NK cells consistently exhibited reduced degranulation (**Fig. 4A, B**), indicating that GABA had a negative impact on NK cell-mediated cytotoxicity in the presence of YAC-1 cells. Next, we tested if GABA impacted the cytotoxicity of NK-92 cells, a human NK cell line. Similar to the experiments with the mouse NK cells, GABA inhibited NK-92 cell-mediated killing of K-562 cells (**Fig. 4C**), confirming that GABA inhibits NK cell cytotoxicity *in vitro*. Finally, we tested GABA-containing supernatant from T. gondii-challenged DCs. While there was a non-significant difference between mouse IL-2-stimulated NK cells pre-treated with supernatant from infected DCs compared with control supernatant, the overall trend was that the supernatants from infected DCs had a negative impact on degranulation by NK cells in the presence of YAC-1 cells (**Fig. 4D, E**). Jointly, the data indicate that GABA reduces NK cell cytotoxicity *in vitro*.

**Figure 4.**
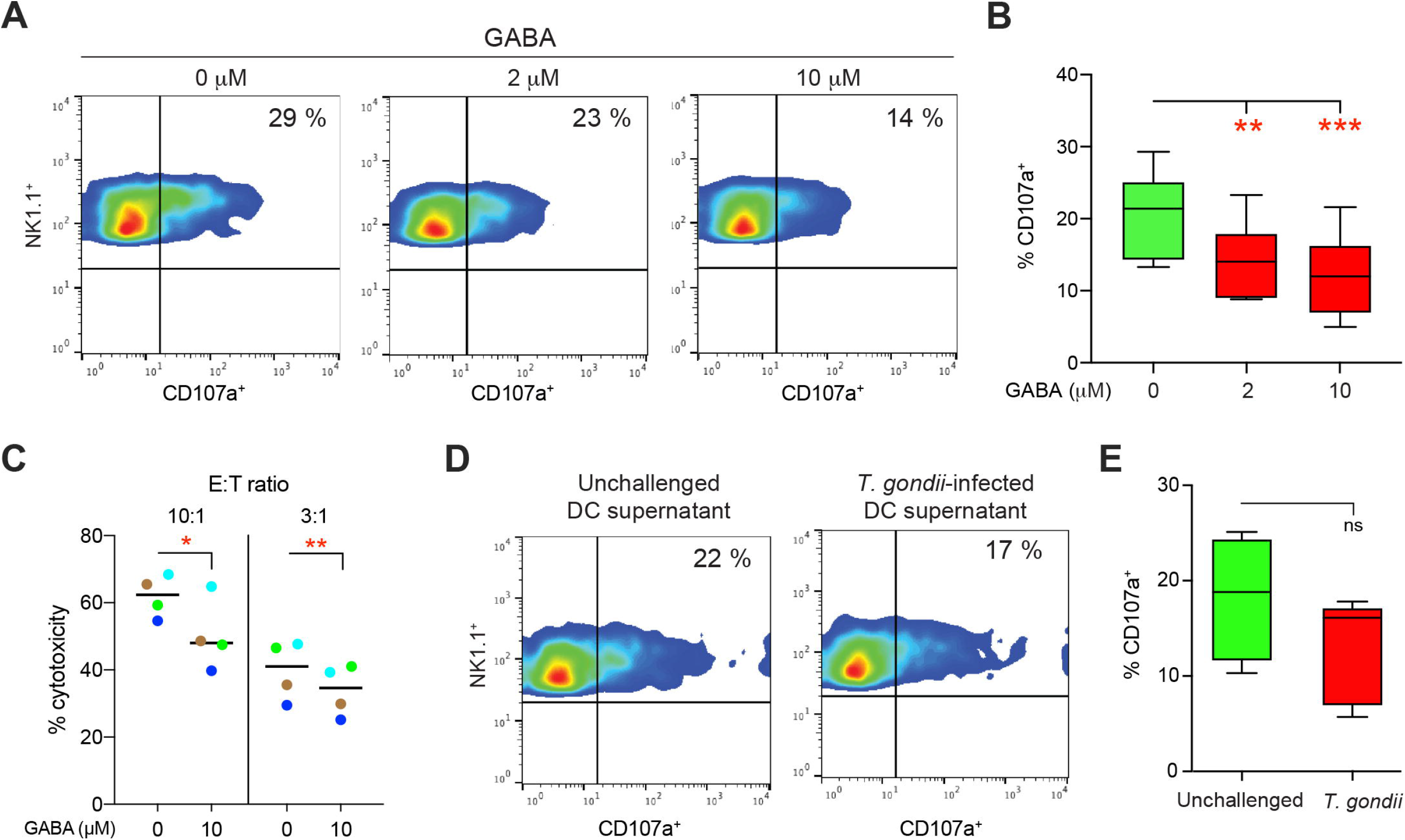
Degranulation of mouse NK cells and cytotoxicity of NK-92 cells in the presence of GABA. **(A-B)** NK cells were preincubated with GABA at indicated concentration for 2 h prior to incubation with YAC-1 cells. (**A**) Representative bivariate plots show % of cells with surface expression of NK1.1 and CD107a. (**B**) Box-and-whisker plots show median values from 6 independent experiments. ** p < 0.01, *** p < 0.001, RM one-way ANOVA followed by Dunnett’s multiple comparison test, (n= 6 independent experiments). (**C**) NK-92 cells were incubated in the presence of GABA for 2 h prior to incubation with ^51^Cr labelled K-562 cells. Graph represents % cytotoxicity calculated as indicated under Methods, * p < 0.05, ** p < 0.01, paired Student’s t-test, (n = 4 independent experiments with 3 technical replicates each). (**D, E**) NK cells were incubated with supernatants from unchallenged mBMDCs or *T. gondii* (RH-LDM)-infected mBMDCs for 2 h prior to incubation with YAC-1 cells for measuring degranulation, as in (A-B). ns: p > 0.05, paired Student’s t-test, (n = 4 independent experiments).

### 3.5 Supernatants from *T. gondii*-challenged NK cells rescue hypermotility in GABA-inhibited parasitized DCs

GABAergic signaling drives the migratory activation of parasitized DCs, termed hypermotility ^3^. We previously reported that transfer of *T. gondii* from parasitized DCs to effector NK cells leads to productive infection of NK cells ^20^. However, whether NK-DC interactions can be modulated by GABAergic signaling has not been addressed. We therefore sought to determine if supernatants from infected NK cells could modulate the motility of parasitized DCs. First, DCs where subjected to GABAergic inhibition with the GABA synthesis inhibitor SC, which abrogated hypermotility of infected DCs, with a non-significant impact on base-line motility of unchallenged D**Cs** (**Fig. 5A, B**). Importantly, GABA-containing supernatant collected from parasitized NK cells (**Fig. 3D**), restored hypermotility of parasitized DCs in presence of SC (**Fig 5A, B**). This reconstitution was abolished by the broad GABA-A R inhibitor picrotoxin, suggesting an indispensable role for GABA-A Rs in hypermotility of DCs. Further, supernatant from NK cells pretreated with SC (**Fig. 3C**) failed to reconstitute hypermotility. Finally, addition of exogenous GABA reconstituted hypermotility in presence of SC. We conclude that GABA-containing NK cell supernatants rescue hypermotility of parasitized DCs. Consistent with the GABAergic signaling cascade ^36^, GABA-containing NK cell supernatants rescued hypermotility when upstream GABA synthesis was inhibited (SC) but not when downstream GABA-A R activation was inhibited (picrotoxin). Thus, GABA-containing supernatants from NK cells impacted the motility of parasitized DCs in a picrotoxin inhibitable fashion which was rescued by exogenous GABA, jointly indicating implication of GABA-A Rs.

**Figure 5.**
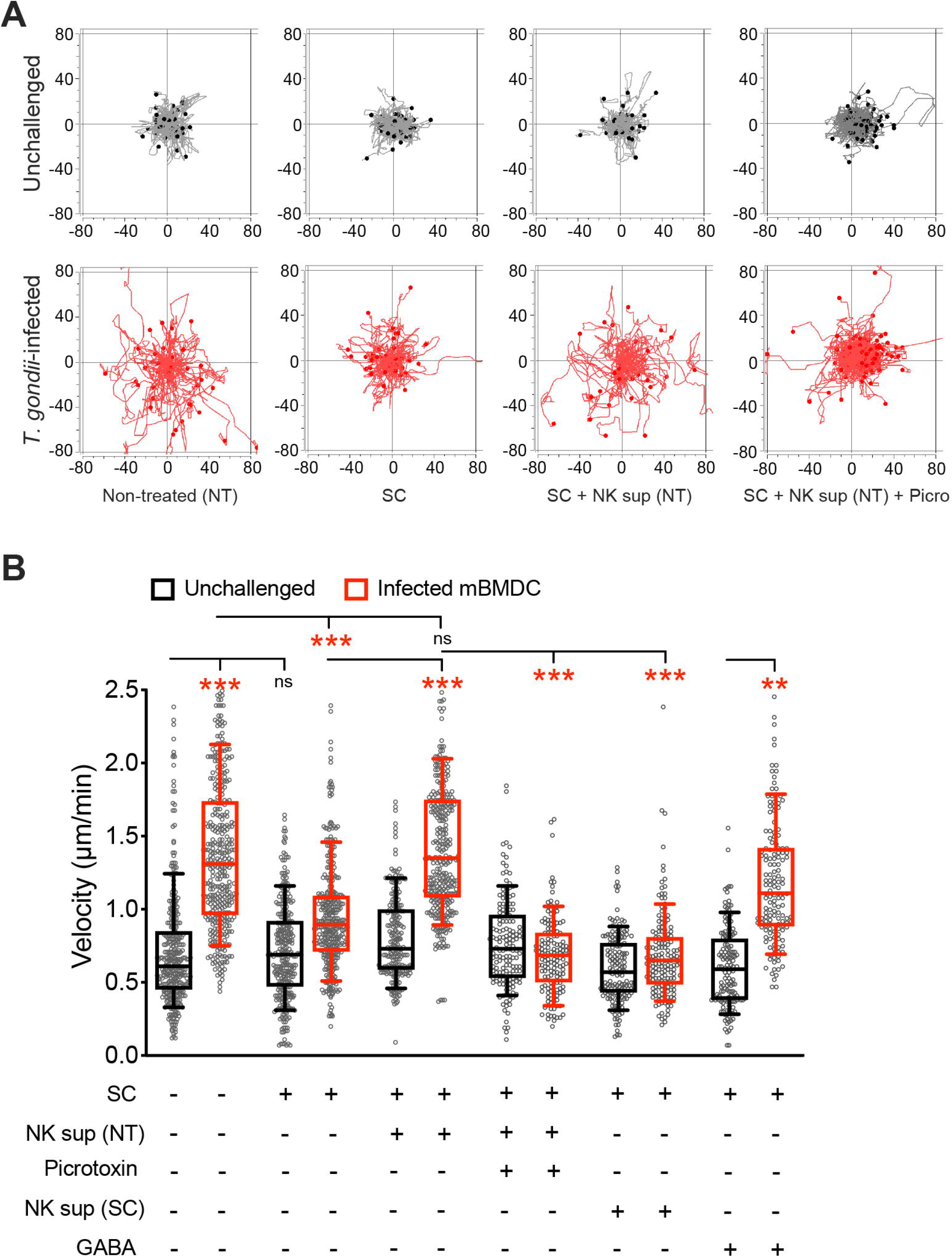
Supernatants from infected NK cells induce hypermotility in mDCs. **(A)** Representative motility plots of unchallenged and *T. gondii* (ME49/PTG)-infected mBMDCs treated with SC in presence and absence of supernatants from T. gondii-infected mNK cells with or without picrotoxin. X-and y-axes show distances in μm. **(B)** Box-and-whisker dot plots show cell velocities (μm/min) for each condition, as in (A). Cells were treated with SC in presence of supernatants from T. gondii-infected mNK cells, picrotoxin or GABA. Non-treated (NT) and SC-treated (SC) mNK cell supernatants were collected as indicated under Methods and added at a 1:1 ratio with medium. ** p < 0.01, *** p < 0.001, ns: p > 0.05, one-way ANOVA followed by Tukey multiple comparison test (n = 3-6 independent experiments per condition).

## 4 Discussion

To date, the GABAergic potential of NK cells has remained unexplored ^11^. Here, we report that NK cells are in fact GABAergic cells and that GABA signaling is implicated in the responses of NK cells challenged with *T. gondii*. Our study establishes that both human and murine NK cells transcriptionally express components of the GABAergic system including GABA synthesis and degradation enzymes, GABA transporters, GABA-A R subunits and regulators (CCCs). However, the overall expression repertoire of GABA-A R subunits was more restricted in NK cells compared with primary microglia ^34^ and more similar to that of DCs ^3^. Commonly expressed GABA-A R subunits for human and mouse NK cells were α3, β2, γ1 and ρ1/2. This expression repertoire in NK cells provides a base for the formation of both heteropentameric (2 α:s + 2 β:s + 1 additional subunit) and homopentameric (ρ:s) GABA-A Rs. Further, in human NK cells, variability in the expressed subunits was observed among donors. Yet, the α3, β2 and ρ2 subunits were commonly expressed by all tested donors. Jointly, this indicates a conserved expression of α, β and ρ subunits for human and murine NK cells.

Cl^-^ transporters (CCCs) regulate the level of intracellular Cl^-^ and ultimately govern the effects of GABA-A R activation ^6^. The expression pattern of CCCs in NK cells was consistent with that found in peripheral blood mononuclear cells ^12^ and microglia ^34^. The CCCs NKCC1 and KCC2 are major regulators of intracellular Cl^-^concentrations in neuronal cells ^37^. Because expression of KCC2 was low or absent in NK cells, NKCC1 is likely the principal determinant for setting the intracellular Cl^-^ concentration and thus, for regulating the effects mediated by GABA-A R activation.

In GABAergic mammalian cells, GABA is chiefly synthesized from glutamate by the action of the enzymes GAD65 and GAD67 ^7^. GAD65 synthesizes GABA mainly for vesicular release and GAD67 for cytosolic release. The expression of GAD67 by peripheral immune cells has remained undetermined. Here, we show that murine NK cells transcribe both enzymes, similar to murine microglia ^34^. In contrast, our data indicate that human NK cells exclusively transcribe GAD67 and express GAD67 protein upon *T. gondii* challenge. Initially, murine T cells, macrophages and DCs were reported to express GAD65 and secrete measurable amounts of GABA ^3,13^. However, a more recent report characterized the expression of GAD65/67 in murine DCs, while human monocytes and DCs exclusively expressed GAD67 ^38^. Thus, expression of GAD67 over GAD65 seems to distinguish human NK cells, and human mononuclear phagocytes, from other peripheral immune cells. Regardless, the data demonstrate abundant release of GABA by both human and murine NK cells upon challenge with *T. gondii*. The upregulation of GAD transcripts likely explains the elevated GABA secretion upon infection, which was inhibited by GAD inhibition. Jointly, this imparts functional roles for GADs in NK cells.

Related to transportation of GABA, only transcripts of GAT2 were detected in both human and murine NK cells. In contrast, GAT2 and GAT4 were expressed by murine microglia and DCs ^3,34^ and GAT1 was dysregulated in T cells in models of multiple sclerosis and experimental autoimmune encephalomyelitis, impacting proliferative and cytotoxic immune responses ^13,39^. Our assumption that GAT2 transports GABA out of NK cells for autocrine stimulation of GABA A Rs is consistent with attributed functions of efflux of GABA by GAT2 in neuronal models, including release of GABA across the blood-brain barrier ^40^. Taken together, the expression of GABAergic signaling components in NK cells and the implication of the GABAergic system of T cells and phagocytes in immunomodulation and responses to infection ^41^ motivated a functional assessment of GABAergic signaling in NK cells.

We show that GABAergic activation has functional implications for NK cell effector functions and interactions with DCs. Importantly, NK cells responded to *T. gondii* infection with GABA secretion. Jointly with GAD67, the expression of GAT2 supports the notion that GABA is synthesized cytosolically and secreted in vesicle-independent fashion for tonic modulations of GABA-A Rs in NK cells, similar to neurons ^40,42^. However, exactly how the elevated GABA synthesis and secretion is orchestrated in NK cells remains to be investigated. Nevertheless, the consistent reciprocal upregulation of the GABA synthesis enzyme and downregulation of GABA-degrading GABA-T in distinct human donors is indicative that GABA production is tightly regulated in NK cells. In DCs and primary microglia, GABA production requires live intracellular *T. gondii* ^3,34^. Similarly, NK cells responded with transcriptional modulation of several components of the GABAergic system upon infection by T. gondii. Cell sorting revealed that GABA was chiefly secreted by infected (GFP^+^) NK cells, corroborating the association to intracellular localization of live parasites. However, a modest but significant by-stander effect was consistently detected in non-infected (GFP^-^) NK cells, which could not be achieved by challenge with heat-inactivated parasites, lysates, supernatants or IL12/IL18 activation. We speculate that this by-stander effect may be mediated by injection of parasite effectors in host cells in absence of invasion ^43^. Altogether, secretion of GABA, upregulation of GABA transporters, upregulation of GABA-A R subunits and GABA signaling regulators CCCs represent an enhanced GABAergic signaling in NK cells upon infection.

We report that GABA downmodulates cytotoxicity and degranulation of NK cells *in vitro*. Previous work has shown that NK cells target parasitized DCs and that cytotoxic attack can lead to infection of NK cells ^20^. Interestingly, cytokine responses, and specifically IFN-γ responses, are downmodulated in parasitized NK cells by unknown mechanisms ^30^. Previous studies have shown that GABA can regulate cytokine release by peripheral blood mononuclear cells and T cells ^44^, and T cell cytotoxicity ^45^ but it has remained unclear what effects GABA has on NK cells ^46^. Here, we show that GABA, which is secreted by parasitized NK cells and DCs, hampers cytotoxicity and degranulation of NK cells *in vitro*. Additionally, secreted GABA also modulated the migratory responses of DCs, shown by the reconstitution of hypermotility in parasitized GABA-inhibited DCs by GABA-containing supernatants from parasitized NK cells or exogenous GABA, but not upon GABA-A R antagonism. Altogether, we speculate that GABA may have dual effects during infection: down-modulation of pro-inflammatory responses and enhancement of DC migration. Hypothetically, in the context of infection in the microenvironment in tissues, this dual effect might facilitate transmission of the parasite between leukocytes and thus facilitate dissemination. However, in contrast to DCs, the migratory responses of parasitized NKs *in vitro* seem to be non-significantly affected ^47^. Because both DCs and NK cells secrete GABA upon *T. gondii* infection, the different migratory responses of the two cell types indicate differences in GABA-A R function. This is also consistent with the present findings that DCs and NKs express different subsets of GABA-A R subunits.

To our knowledge, this represents the first report of GABAergic signaling in NK cells. The data provide a proof-of-concept that GABAergic signaling can modulate NK cell functions in the context of infection. We have recently shown that *T. gondii* infection impacts MAP kinase activation and calcium signaling in immune cells ^4,36,48^. Thus, immunomodulatory effects in parasitized NK, for example cytokine secretion, could be mediated through MAP kinase signaling. If GABAergic signaling acts on NK cells via voltage-gated calcium channels awaits further investigation. Taking into account recent reports showing that T cells, DCs, monocytes, macrophages and microglia are GABAergic cells ^34,36,44^, the mounting evidence advocates that GABAergic signaling may be conserved throughout the immune system. GABAergic activation in immune cells is likely not restricted to intracellular parasitism and, hypothetically, encompasses additional immune cell functions.

## Authorship

AKB, LMF, SK, SJ-B, IE-L, AKW, BJC performed experimental work. AKB, AKW, BJC and AB analyzed the data. AKW and BJC contributed human and murine materials. AKB, BJC and AB prepared figures and wrote manuscript.

## Acknowledgements

This work was funded by the Swedish Research Council (Vetenskapsrådet, 2018-02411 to AB, 2017-00923 to BJC), the Olle Engkvist Foundation (193-609 to AB) and Cancerfonden (CAN 2017/714 to BJC).

## Conflict of interest disclosure

Authors declare that research was conducted in the absence of any commercial or financial relationships that could be construed as a potential conflict of interest.

## Notes

### Competing Interest Statement

The authors have declared no competing interest.

