## supplementary tables and figures for "GABAergic signaling in human and murine NK cells upon challenge with *Toxoplasma gondii*"

**Supplementary Table 1. Primer pair sequences used in real-time quantitative PCR**

| mouse mRNA target | Forward primer sequence | Reverse primer sequence | Amplicon |
|---|---|---|---|
| GABA-A R subunits | | | |
| α1(*GabrA1*) | AAAAGCGTGGTTCCAGAAAA | GCTGGTTGCTGTAGGAGCAT | 84 |
| α2 (*GabrA2*) | GCTACGCTTACACAACCTCAGA | GACTGGCCCAGCAAATCATACT | 115 |
| α3 (*GabrA3*) | GCCGTCTGTTATGCCTTTGTATTT | TTCTTCATCTCCAGGGCCTCT | 119 |
| α4 (*GabrA4*) | AGAACTCAAAGGACGAGAAATTGT | TTCACTTCTGTAACAGGACCCC | 118 |
| α5 (*GabrA5*) | GATTGTGTTCCCCATCTTGTTTGGC | TTACTTTGGAGAGGTGGCCCCTTTT | 100 |
| α6 (*GabrA6*) | TGGGAGCTATGCTTATCCT | TCTTCTGGGACTTCTACTGAG | 81 |
| β1 (*GabrB1*) | GGTTTGTTGTGCACACAGCTCC | ATGCTGGCGACATCGATCCGC | 153 |
| β2 (*GabrB2*) | GCTGGTGAGGAAATCTCGGTCCC | CATGCGCACGGCGTACCAAA | 70 |
| β3 (*GabrB3*) | GAGCGTAAACGACCCCGGGAA | GGGACCCCCGAAGTCGGGTCT | 100 |
| γ1 (*GabrG1*) | ATCCACTCTCATTCCCATGAACAGC | ACAGAAAAAGCTAGTACAGTCTTTGC | 100 |
| γ2 (*GabrG2*) | ACTTCTGGTGACTATGTGGTGAT | GGCAGGAACAGCATCCTTATTG | 147 |
| γ3 (*GabrG3*) | ATTACATCCAGATTCCACAAGATG | CAC AGG TGT CCT CAA ATT CCT | 149 |
| δ (*GabrD*) | GAATCCGTTCCAGACTCAAA | GCACTAGGCTCAACTTCAGG | 349 |
| ε (*GabrE*) | ACTGCGCCCTGGCATTGGAG | AGGCCCGAGGCTGTTGACAA | 70 |
| θ (*GabrQ*) | GCTGGAGGTGGAGAGCTATGGCT | CCCCAGGTACGTGTACTGAGGGA | 115 |
| π (*GabrP*) | CAGAGGACGTGCATCCAGGGGA | TCCGAACTGGGTCACCACCGAA | 139 |
| ρ1 (*GabrR1*) | CTGGAAATCGAAAGCTACGC | AGATGTGACGACGCAGAGTG | 205 |
| ρ2 (*GabrR2*) | CCAAGCCAAGCCATTTGTAT | GTCCCTCCAGTAATGCCTCA | 227 |
| ρ3 (*GabrR3*) | CAACTCAACAGGAGGGGAAA | TCCACATCAGTCTCGCTGTC | 101 |
| Enzymes | | | |
| GAD65 (*Gad2*) | GCTGGAACCACCGTGTATGG | TCCACGTGCATCCAGATCTTAT | 86 |
| GAD67 (*Gad1*) | GTGACCAGGGTGCCCGCTTC | TGCGCAGTTTGCTCCTCCCC | 100 |
| GABA-T (*Abat*) | TGGCCTTCTTGTTGATTACC | TATATCTGGATCCAGGTATGAAGAG | 81 |
| Transporters | | | |
| GAT1 (*Slc6a1*) | TGAACTCTTCATTGCTGCC | AAGACATAAATGCCACCCTG | 81 |
| GAT2 (*Slc6a12*) | CATGATGCCTTTGTCCCAG | TACAGACGAACTGGCTGTC | 83 |
| GAT3 (*Slc6a13*) | ATCTTTGAAGGCATCGGCT | GAAGAGGTAGAAGAGGG | 96 |
| GAT4 (*Slc6a11*) | Mm_Gabt4_1_SG QuantiTect Primer Assay (QT00130116, Quiagen) | | 78 |

| | | | |
|---|---|---|---|
| CCCs | | | |
| NKCC1 (*Slc12a2*) | GATGCTGTGGTCGCATACACT | CAGCGGACTAATACACCCTTG | 77 |
| NKCC2 (*Slc12a1*) | CTGGCTAAGAATGTGACTGT | TCTGCTTGCTCATCTCCAG | 83 |
| KCC1 (*Slc12a4*) | TCTACCTGGGGACGACATTTG | CCGATGGGTAAAAGATGGCAG | 102 |
| KCC2 (*Slc12a5*) | TCAGTCACAGGGATCATGG | GGATAGTTCCAGTAGGGATAGAC | 82 |
| KCC3 (*Slc12a6*) | CTGCCATCTTTCGGAGTGACG | AGAAGGCTGTACCATAGACGC | 81 |
| KCC4 (*Slc12a7*) | ATGCCCACGAACTTTACGGTG | GGGAGGTTTGATCCACGCT | 115 |
| Reference genes | | | |
| TATA-binding protein (*TBP*) | GGGGAGCTGTGATGTGAAGT | CCAGGAAATAATTCTGGCTCA | 93 |
| Importin 8 (*IPO8*) | CTATGCTCTCGTTCAGTATGC | GTCCGAAAGATCTCCATCCA | 81 |

| **human mRNA target** | **Forward primer sequence** | **Reverse primer sequence** | **Amplicon** |
|---|---|---|---|
| GABA-A R subunits | | | |
| α1(*GabrA1*) | GTCACCAGTTTCGGACCCG | AACCGGAGGACTGTCATAGGT | 119 |
| α2 (*GabrA2*) | GTTCAAGCTGAATGCCCAAT | ACCTAGAGCCATCAGGAGCA | 160 |
| α3 (*GabrA3*) | CAACTTGTTTCAGTTCATTCATCCTT | CTTGTTTGTGTGATTATCATCTTCTTAGG | 102 |
| α4 (*GabrA4*) | TTGGGGGTCCTGTTACAGAAG | TCTGCCTGAAGAACACATCCA | 105 |
| α5 (*GabrA5*) | TTGGATGGCTACGACAACAGA | GTCCTCACCTGAGTGATGCG | 62 |
| α6 (*GabrA6*) | ACCCACAGTGACAATATCAAAAGC | GGAGTCAGGATGCAAAACAATCT | 67 |
| β1 (*GabrB1*) | TGCATGTATGATGGATCTTCG | GTGGTATAGCCATAACTTTCGA | 80 |
| β2 (*GabrB2*) | GCAGAGTGTCAATGACCCTAGT | TGGCAATGTCAATGTTCATCCC | 137 |
| β3 (*GabrB3*) | CAAGCTGTTGAAAGGCTACGA | ACTTCGGAAACCATGTCGATG | 108 |
| γ1 (*GabrG1*) | CCTTTTCTTCTGCGGAGTCAA | CATCTGCCTTATCAACACAGTTTCC | 91 |
| γ2 (*GabrG2*) | CACAGAAAATGACGGTGTGG | TCACCCTCAGGAACTTTTGG | 136 |
| γ3 (*GabrG3*) | AACCAACCACCACGAAGAAGA | CCTCATGTCCAGGAGGGAAT | 113 |
| δ (*GabrD*) | CTTTGCTCATTTCAACGCC | TTCCTCACGTCCATCTCTG | 86 |
| ε (*GabrE*) | ACAGGAGTGAGCAACAAAACTG | TGAAAGGCAACATAGCCAAA | 107 |
| θ (*GabrQ*) | CCAGGGTGACAATTGGCTTAA | CCCGCAGATGTGAGTCGAT | 63 |
| π (*GabrP*) | CAATTTTGGTGGAGAACCCG | GCTGTCGGAGGTATATGGTG | 110 |
| ρ1 (*GabrR1*) | GAAAGGCAGCCCAATTCTG | TCTATCCTCAGAAGCTGTTCTG | 80 |
| ρ2 (*GabrR2*) | TACAGCATGAGGATTACGGT | CAAAGAACAGGTCTGGGAG | 81 |
| ρ3 (*GabrR3*) | TGATGCTTTCATGGGTTTCA | CGCTCACAGCAGTGATGATT | 111 |
| Enzymes | | | |
| GAD65 (*Gad2*) | CTACGCGTTTCTCCATGCAA | GCCAAAGTGGGCCTTTCTC | 63 |
| GAD67 (*Gad1*) | GCGGACCCCAATACCACTAAC | CACAAGGCGACTCTTCTCTTC | 144 |
| GABA-T (*Abat*) | TGAAATACCCTCTGGAAGAG | CAATCAGATCCTCCACCTC | 80 |
| Transporters | | | |
| GAT1 (*Slc6a1*) | TTTCTGTGCCTGTTACCCA | ATGTCTCGGAATTTGGACG | 126 |

| | | | |
|---|---|---|---|
| GAT2 (*Slc6a13*) | CTTCACTAAGGTGGGATGGA | TTCCATGACTGGATACACTGG | 81 |
| GAT3 (*Slc6a11*) | TATTTGAAGGCATTGGCTATGC | GTAGTGAAGCAGTTGCTCAG | 108 |
| CCCs | | | |
| NKCC1 (*Slc12a2*) | TGGGTCAAGCTGGAATAGGTC | ACCAAATTCTGGCCCTAGACTT | 161 |
| NKCC2 (*Slc12a1*) | TCAGGAGATTTGGAGGATCCC | ACCCCTAAGTAGGCAACAGTG | 86 |
| KCC1 (*Slc12a4*) | CCTCCCGTGTTTCCGGTATG | CAGGAGTCGGTCGTAAGGTTG | 155 |
| KCC2 (*Slc12a5*) | GGAAGGAAATGAGACGGTGA | TCCCACTCCTCTCCACAATC | 200 |
| KCC3 (*Slc12a6*) | GGATGTCATCGAGGACCTGAG | TCGAGCTTTCTTATGTCCGTC | 82 |
| KCC4 (*Slc12a7*) | ATCTACTTCCCTTCCGTGACC | TCTGTGCATCCTTGAGGTCC | 70 |
| Reference genes | | | |
| TATA-binding protein (*TBP*) | GAGCTGTGATGTGAAGTTTCC | TCTGGGTTTGATCATTCTGTAG | 117 |
| Importin 8 (*IPO8*) | GCAAAGGAAGGGGAATTGAT | CGAAGCTCACTAGTTTTGACCC | 91 |

Primer pair sequences were designed using GETprime or NCBI primer blast databases.

**Supplementary Table 2**. Modulated transcriptional expression of GABA-A R subunits, enzymes, GABA transporter and CCCs in *T. gondii*-challenged primary mNK cells (related to Figure 2).

| mNK | 4 h | | 24 h | |
|---|---|---|---|---|
| | **Mean** | **SEM** | **Mean** | **SEM** |
| **GABA-A R subunits** | | | | |
| α2 | -91.28 | 5.51 | 57.14 | 20.18 |
| α3 | -58.27 | 9.72 | n.a | - |
| β2 | -94.92 | 3.02 | 98.04 | 1.96 |
| β3 | -62.48 | 31.93 | 89.70 | 10.30 |
| γ1 | 67.06 | 24.16 | 99.33 | 0.67 |
| γ2 | n.a | - | 86.81 | 13.19 |
| δ | -23.06 | 10.74 | -35.72 | 22.54 |
| ε | 43.16 | 51.93 | -20.78 | 18.14 |
| ρ1 | 59.02 | 34.11 | 71.26 | 13.25 |
| ρ2 | 99.55 | 8.36 | 99.50 | 0.50 |
| **Enzymes** | | | | |
| GAD 65 | 170.14 | 69.55 | 87.01 | 20.33 |
| GAD 67 | 142.84 | 24.17 | 677.92 | 48.07 |
| GABA-T | 181.11 | 42.54 | 123.79 | 34.27 |
| **Transporter** | | | | |
| GAT2 | 50.80 | 18.42 | 56.45 | 2.77 |
| **CCCs** | | | | |
| NKCC1 | 2.26 | 1.11 | 17.72 | 21.37 |
| NKCC2 | 97.99 | 2.01 | 4.83 | 2.50 |
| KCC1 | 16.86 | 2.47 | 2.50 | 6.73 |
| KCC2 | -0.87 | 10.91 | -82.54 | 1.87 |
| KCC3 | -14.69 | 0.58 | -24.01 | 11.61 |
| KCC4 | 22.96 | 1.22 | 99.09 | 10.20 |

The relative mRNA expression ($2^{-\Delta Ct}$) was determined as detailed in Materials and Methods. Data show percentage increase or decrease (mean and SEM) of mRNA levels in *T. gondii*-challenged primary mNK cells in comparison to unchallenged cells at indicated time points from 3-5 independent experiments. n.a: not amplified.

**Supplementary Table 3.** Modulated transcriptional expression of GABA-A R subunits, enzymes, GABA transporter and CCCs in *T. gondii*-challenged primary hNK cells (related to Figure 2).

| **hNK** | **24 h** | |
|---|---|---|
| | **Mean** | **SEM** |
| **GABA-A R subunits** | | |
| α3 | 41.20 | 18.46 |
| α5 | 326.89 | 57.41 |
| α6 | 885.23 | 88.02 |
| β2 | 248.57 | 163.57 |
| γ1 | -53.88 | 11.61 |
| δ | 577.50 | 78.15 |
| ρ1 | 74.77 | 79.95 |
| ρ2 | -36.01 | 4.32 |
| **Enzymes** | | |
| GAD 67 | 143.58 | 67.46 |
| GABA-T | -40.14 | 13.18 |
| **Transporter** | | |
| GAT2 | 20.38 | 11.51 |
| **CCCs** | | |
| NKCC1 | 94.20 | 3.15 |
| KCC1 | 53.80 | 8.12 |
| KCC3 | -28.72 | 8.99 |
| KCC4 | -12.70 | 20.03 |

The relative mRNA expression ($2^{-\Delta Ct}$) was determined as detailed in Methods. Data indicate percentage increase or decrease (mean and SEM) of mRNA levels in *T. gondii*-challenged primary hNK cells in comparison to unchallenged cells at indicated time points from 3-4 independent experiments.

**Supplementary Table 4**. Correlation analyses of mRNA expression levels with amounts of GABA released by *T. gondii*-challenged NK cells from individual human donors

| GABA conc. vs | Spearman R | P value |
|---|---|---|
| **GABA-A R subunits** | | |
| α3 | 0.8 | 0.33 |
| α5 | 0 | >0.99 |
| α6 | -0.2 | 0.92 |
| β2 | 0.2 | 0.92 |
| γ1 | 0.6 | 0.42 |
| δ | -0.5 | >0.99 |
| ρ1 | 0.8 | 0.33 |
| ρ2 | 0.4 | 0.75 |
| **GABA enzymes** | | |
| GAD 67 | 0.95 | 0.0004 |
| GABA-T | -0.87 | 0.0045 |
| **GABA transporter** | | |
| GAT2 | 0.8 | 0.33 |
| **Chloride co-transporters (CCC)** | | |
| NKCC1 | 0.6 | 0.42 |
| KCC1 | 0.4 | 0.75 |
| KCC3 | -0.8 | 0.33 |
| KCC4 | 0 | >0.99 |

Correlations were assessed by Spearman test (related to Figure 3).

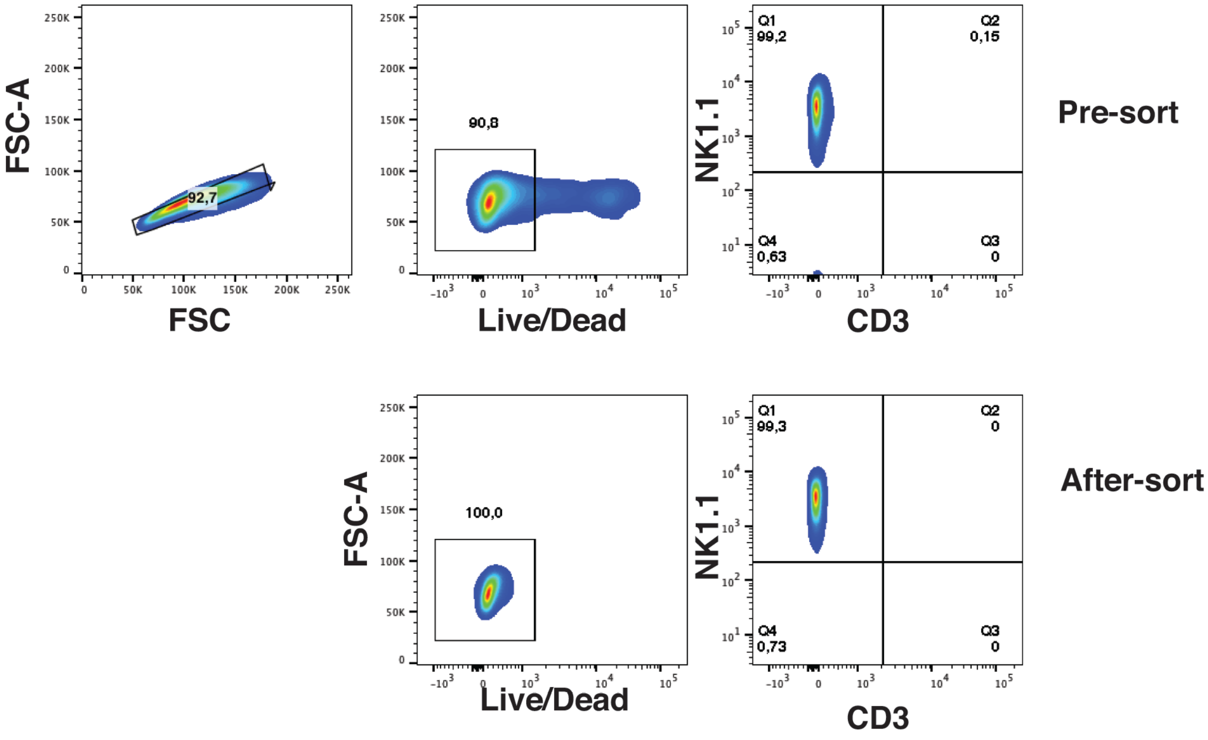


**Supplementary Figure 1.** Representative flow cytometric analysis of cell purity of mNK cells used for mRNA analyses of GABA-related genes

Cells were first isolated by negative sorting using NK cell isolation kit (Pre-sort) and further purified by flow cell sorting (After-sort) following labelling with anti-CD3, anti-NK1.1 and Live/Dead staining, as detailed under Methods.

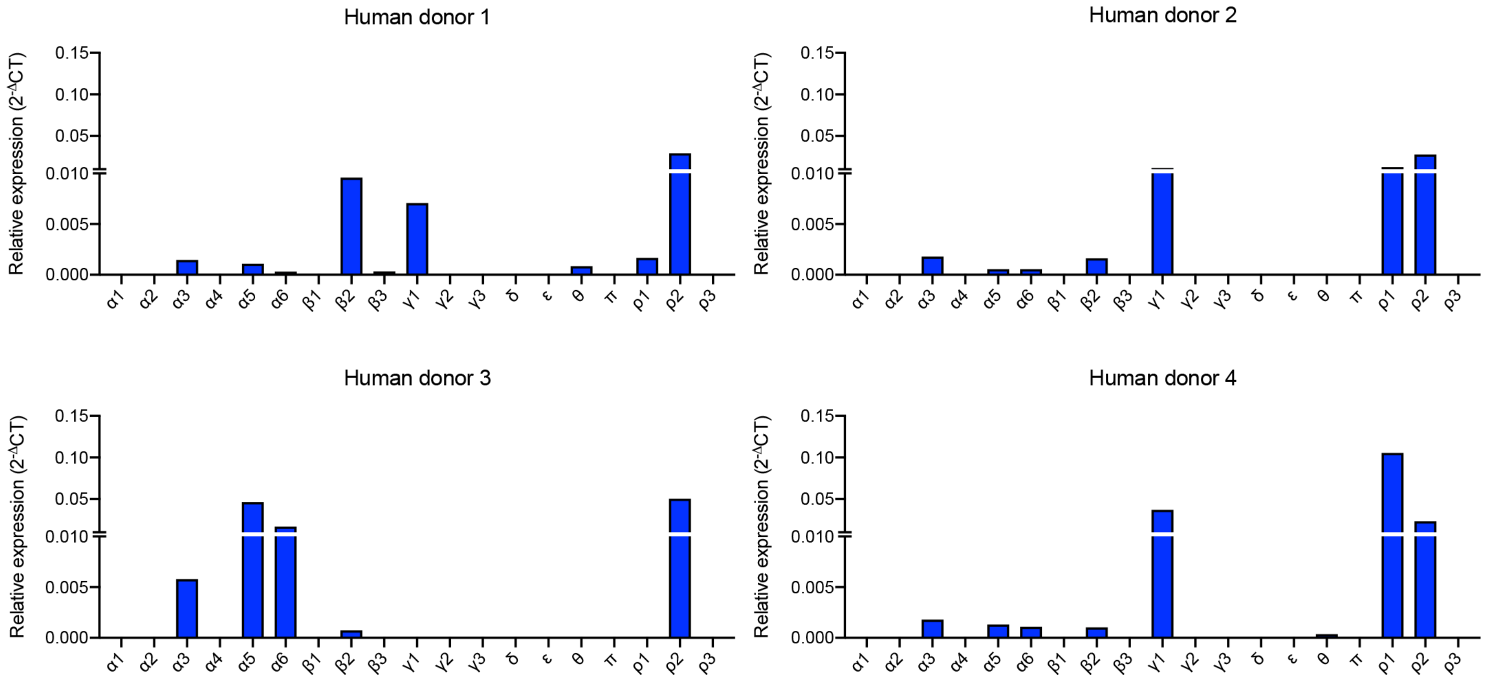


**Supplementary Figure 2.** Relative mRNA expression of 19 GABA-A R subunits in NK cells from four different human donors (donor 1-4). NK cells were purified from human blood and qPCR was performed as indicated under Methods.
